# Genetic diversity and population history of eight Italian beef cattle breeds using measures of autozygosity

**DOI:** 10.1101/2021.02.22.432240

**Authors:** Maria Chiara Fabbri, Christos Dadousis, Francesco Tiezzi, Christian Maltecca, Emmanuel Lozada-Soto, Stefano Biffani, Riccardo Bozzi

**Affiliations:** Dipartimento di Scienze e Tecnologie Agrarie, Alimentari, Ambientali e Forestali, Università di Firenze, Firenze, Italy; Dipartimento di Scienze Medico-Veterinarie, Università di Parma, Parma, Italy; Department of Animal Science, North Carolina State University, Raleigh, NC 27695, United States; Institute of Agricultural Biology and Biotechnology (CNR), Milano, Italy

## Abstract

In the present study, GeneSeek GGP-LDv4 33k single nucleotide polymorphism chip was used to detect runs of homozygosity (ROH) in eight Italian beef cattle breeds; six breeds with distribution limited to Tuscany (Calvana, Mucca Pisana, Pontremolese) or Sardinia (Sarda, Sardo Bruna and Sardo Modicana) and two cosmopolitan breeds (Charolais and Limousine). ROH detection analyses were used to estimate autozygosity and inbreeding and to identify genomic regions with high frequency of ROH, which might reflect selection signatures. Comparative analysis among breeds revealed differences in length and distribution of ROH and inbreeding levels. The Charolais, Limousine, Sarda, and Sardo Bruna breeds were found to have a high frequency of short ROH (30.000); Calvana and Mucca Pisana presented also runs longer than 16 Mbp. The highest level of average genomic inbreeding was observed in Tuscan breeds, around 0.3, while Sardinian and cosmopolitan breeds showed values around 0.2. The population structure and genetic distances were analyzed through principal component analysis and multidimensional scaling analysis, and resulted in a clear separation among the breeds, with clusters related to productive purposes and sample sizes. The frequency of ROH occurrence revealed eight breed-specific genomic regions where genes of potential selective and conservative interest are located (e.g. *MYOG, Chitinases* (BTA16), *TIMELESS*, *APOF*, Olfactory receptors, *CACNG2* (BTA5) and Collagens (BTA2)). In all breeds, we found the largest proportion of homozygous by descent segments to be those that represent inbreeding events that occurred around 32 generations ago; with Tuscan breeds also having a significant proportion of segments relating to more recent inbreeding.

## Introduction

Runs of homozygosity (ROH) consist of contiguous regions of the genome where an individual is homozygous in all sites [1]. This occurs when the haplotypes transmitted from both parents are identical due to being inherited from a common ancestor. The length of a ROH is an imprint of the history of a population linked to its effective population size and provides evidence for phenomena such as inbreeding, mating system, and population bottlenecks. In theory, longer ROH are due to recent inbreeding, as recombination has not had the possibility of breaking up the homozygous segment, on the other hand, short ROH demonstrate an older origin because several meiosis have been occurred [2]. Information on inbreeding is crucial in the design of breeding and conservation programs to control the increase in inbreeding levels and to avoid the unfavorable effect of inbreeding depression in progeny [3].

The inbreeding coefficient of an individual (F) is defined as the probability that two randomly chosen alleles at a specific locus within an individual are identical by descent (IBD) [4]. Homozygosity caused by two IBD genomic segments is defined “autozygosity”, F is therefore an estimate of genome-wide autozygosity [5] and ROH are highly likely to be autozygous [6].

The estimation of the inbreeding coefficient from the proportion of the genome covered by ROH (F_ROH_) has been considered a powerful and accurate method of detecting inbreeding effects [5] and a valid alternative to pedigree inbreeding coefficient [7, 8], which doesn’t take into account the stochastic nature of recombination; pedigree information could be incomplete and/or incorrect especially for local breeds, where the extensive breeding system and the natural mating system could allow a limited control of relatedness.

The high occurrence of ROH in chromosomes could potentially represent a selection signature, i.e. a genomic footprint that could provide an overview for understanding the mechanism of selection and adaptation, and could help to uncover regions related to important physiological, economical and adaptive traits [10]. A selection signature is characterized by a reduced haplotype variability, defined as ROH island [11]. Two different methods have been applied to detect ROH islands: the first one based on an arbitrarily defined frequency of common ROH within the population (for e.g., 20 % [12]; 45% [13]; 70% [14]), while the second approach on a percentile threshold (99th percentile) based on the top 1% of SNPs observed in a ROH [15, 16].

However, the use of ROH as markers for the identification of genomic regions potentially subjected to non-recent evolutionary events is not straightforward. It requires that homozygous segments have been inherited from old ancestors and were not caused by recent demographic events [17]. A further approach to estimate global inbreeding (F_G_) for each population, which links the genomic homozygous segments to the time of living of the most recent common ancestor, is the Homozygous-Identical by Descent (HBD) state probabilities. Druet and Gautier [18] presented an approach to investigate local and global inbreeding, based on the hidden Markov model (HMM). This approach assumes that the genome is formed by HBD and non-HBD segments, where each segment has a HBD state probability. Solé et al. [19] implemented a new HMM with multiple age based HBD-classes in which the length of HBD segments have distinct distributions: longer segments for more recent common ancestors, and shorter for more ancient ancestors. The expected HBD segment lengths are inversely related to the number of generations to the common ancestor and their frequency to past effective population size and individual inbreeding coefficients [18].

Assessment of genetic diversity and population structure is an important task to understand the evolutionary history of the breeds, but also to provide important information for the conservation and management of biodiversity [20]. Italy has a biodiversity reservoir for local breeds, but generally, local populations have a small sample size and one of the most important obstacle is the increase in inbreeding, leading to negative effect on production and reproduction traits [21]. The maintenance of genetic diversity should be the priority for countries such as Italy, where local breeds guarantee the economical survivor of marginal areas. Selection programs are not easy to apply to local populations for the reduced sample size which also implies a higher level of inbreeding than in selected breeds [22]. Nevertheless, should be necessary to provide conservation programs maintaining genetic diversity, and to find strategies to control inbreeding levels. Within this context, improving the knowledge about the genomic background of local breeds is crucial.

The aim of this study was to assess genome-wide autozygosity in six Italian beef breeds at critical risk of extinction, mainly reared in Tuscany and Sardinia, namely Calvana (CAL), Mucca Pisana (MUP), Pontremolese (PON), Sardo Bruna (SAB), Sardo Modicana (SAM), Sarda (SAR) and the cosmopolitan Charolais (CHA) and Limousine (LIM), by : i) investigating ROH distribution and characterization across the genome, and consequently, the inbreeding coefficients (F_ROH_) within each breeds; ii) using the HDB state probabilities to calculate global inbreeding (F_G_) and investigate its change across generations, in order to describe the demographic history of the populations.

## Materials and methods

### Ethics statement

DNA sampling for all the eight breeds was conducted using nasal swabs and no invasive procedures were applied. Thus, in accordance to the 2010/63/EU guide and the adoption of the Law D.L. 04/03/2014, n.26 by the Italian Government, an ethical approval was not required for our study.

### Animal sampling, quality control and SNPs characterization

A total of 1,308 animals, belonging to eight breeds, have been genotyped with GeneSeek GGP-LDv4 33k (Illumina Inc., San Diego, California, USA) single nucleotide polymorphism (SNP) DNA chip. Sampled animals for the three Tuscan breeds were 179, 190 and 45 for CAL, MUP and PON, respectively. The limited number of alive animals of these breeds restricted the total samples. Also, for SAM, being at risk of extinction, only 101 genotypes have been recovered. Samples of SAR (n=199) and SAB (n=194) were animals born from 2005 and 2000, respectively. CHA and LIM samples consisted of cattle born from 2015 to 2019 (200 samples for each breed). Genotype quality control and data filtering were performed with PLINK v1.9 [23] and were conducted separately for each breed: only SNPs located on the 29 autosomes were included (n = 28,289). Linkage Disequilibrium (LD) pruning was not performed as suggested by Dixit et al. [24], as LD is related to various evolutionary forces which are the phenomena ROH analysis investigates (e.g. inbreeding, nonrandom mating, population bottleneck, artificial and natural selection). Editing for SNP MAF was not applied to the dataset because it does not improve ROH detection, on the contrary homozygous regions could be ignored [25]. A SNP characterization was performed based on MAF categories. SNPs were classified into 5 classes: monomorphic SNPs, SNPs with MAF ranged from 0 to 0.005, from 0.005 to 0.01, from 0.01 to 0.05 and >0.05. The aim was to evaluate the number of monomorphic SNPs within and between breeds, given that numerous common monomorphic SNPs could influence ROH investigation.

### Multidimensional scaling plot analysis

A multidimensional scaling plot analysis (MDS) was performed to investigate the population structure between the eight breeds based on genetic distances. The first three dimensions were obtained with PLINK v1.9 [23] using the *--mds-plot* flag and calculated on the matrix of genome wide pairwise Identical by State (IBS) distances [26]. Results were plotted using Scatterplot3d R package [27]

### Runs of homozygosity detection

Analysis of runs of homozygosity was conducted with the R package *detectRUNS* v. 0.9.5 [28], using the consecutive method, to avoid the detection of artificial ROH shorter than the window chosen [29]. The following parameters were applied in order to detect a ROH: i) the minimum number of consecutive SNPs was set to 15; ii) the minimum ROH length required was 1 Mbp; iii) the maximum gap between consecutive homozygous SNPs was 1 Mbp; iv) the maximum number of opposite genotypes in the run was set to 2; v) the maximum number of missing genotypes allowed was 2.

A principal component analysis (PCA) was conducted on the number of ROH per chromosome for each breed, to infer the similarities between populations based on ROH chromosomic distribution. All ROH were classified into five classes of length as suggested by Kirin et al. [2], and Ferenčaković et al. [30]: 0-2, 2-4, 4-8, 8-16, >16 Mbp. For each of the eight breeds the total number of ROH, the ROH average number per individuals, the average length of ROH, the number of ROH per breed per chromosome, and the number of ROH per class of length were estimated.

### Genomic inbreeding based on ROH

The genomic inbreeding (F_ROH_) was calculated as suggested by McQuillan et al. [31]:

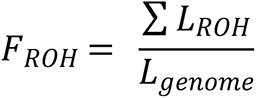

Where ∑ *L_ROH_* was the sum of the length of all ROH found in an individual and *L_genome_* was the total genome length. The F_ROH_ per class of ROH length was calculated.

### Selection signatures and Gene enrichment

In order to investigate the selection signatures in the eight cattle breeds, the occurrences of ROH across genome was explored. The SNPs frequencies (%) in detected ROH were evaluated for each breed and plotted against the position of the SNP across autosomes. The threshold considered was the 80% of ROH occurrence for each breed, which were filtered taking only the genomic regions containing a minimum number of 15 SNPs. These genomic regions were analyzed and overlapped to Genome Data Viewer (https://www.ncbi.nlm.nih.gov/genome/gdv/) of NCBI (National Center for Biotechnology information) to identify genes. The UMD 3.1 assembly was used for mapping. Information on genes detected were obtained from GeneCards database (https://www.genecards.org/).

### Homozygosity by descent (HBD) segments and global inbreeding

The hidden Markov model (HMM)-based approach was used to scan the individual genome for the HBD segments as described in Solé et al. [19]. The analysis was computed with the *RZooROH* R package (https://CRAN.Rproject.org/package=RZooRoH) [32]. The HBD state probability values for each marker were averaged across individual in each population. Averaging HBD probabilities of all loci across the genome led to global (genome-wide) inbreeding (F_G_) calculation. Each class (K) has its own rate parameter, R_K_, which indicates the length of the segments for its respective class. The length of HBD classes is exponentially distributed with rate R_K_, which is double the number of generations to the common ancestor of the respective class. The length of the HBD segment is 1/R_K_ Morgans, indicating high rates associated with shorter segments. The study focused on <16 Mbp ROH length, so the model applied was six HBD classes with respective rates (R_K_ = 2^1^, 2^2^, 2^3^, …, 2^6^) and one non-HBD with an R_K_ rate of 2^6^, so that 32 generations (generation = R/2) and short HBD segments from 1.5 Mbp (1/2^6^) of length were included in the analysis. The rate of the non-HBD class was fixed as the most ancient class.

A MixKR [19] model with K = 7 was performed. To estimate the inbreeding coefficient, we considered the ancestors with an R_K_ rate higher than a threshold T as unrelated. The corresponding genomic inbreeding coefficient (F_G−T_) was then estimated, with R_K_ ≤ T averaged over the whole genome (as reported by Druet et al. [33]).

## Results

### Animal sampling, quality control and SNPs characterization

In total, 28,178 SNPs were divided into 5 classes of MAF (Table 1) while 111 SNPs remained unclassified. PON presented the highest number of monomorphic SNPs (n=4,151; i.e. the 15.6% of the total number of SNPs), followed by LIM and CHA, namely 3,915 (14.4%) and 3,456 (12.8 %) respectively. CAL had an intermediate value (2,968 - 11.40%) while MUP, SAB, SAM and SAR had less than 2,000 monomorphic SNPs, which maintained lower than the 8% of the total amount of them. The first two classes of MAF (0-0.005 and 0.005-0.01) contained few SNPs, exceeding 1,000 markers only in MUP.

**Table 1.** Number of autosomal SNPs per breed^1^ classified into 5 classes of minor allele frequency.

|  | **CAL** | **CHA** | **LIM** | **MUP** | **PON** | **SAB** | **SAM** | **SAR** |
| --- | --- | --- | --- | --- | --- | --- | --- | --- |
| **Monomorphic** | 2,968 | 3,456 | 3,915 | 1,999 | 4,151 | 1,903 | 1,663 | 1,447 |
| **0-0.005** | 589 | 544 | 234 | 912 | 0 | 588 | 313 | 396 |
| **0.005-0.01** | 323 | 123 | 109 | 477 | 0 | 290 | 242 | 253 |
| **0.01-0.05** | 1,359 | 714 | 759 | 1653 | 1628 | 893 | 1,162 | 1,308 |
| **>0.05** | 23,032 | 23,450 | 23,271 | 23,230 | 22,473 | 24,609 | 24,887 | 24,878 |

<sup>1</sup> CAL = Calvana; CHA = Charolais; LIM = Limousine; MUP = Mucca Pisana; PON = Pontremolese; SAB = Sardo Bruna; SAM = Sardo Modicana; SAR = Sarda.

In order to investigate the presence of common SNPs between breeds, pairs comparisons have been performed. Fig 1 explains the total number of monomorphic SNPs on the diagonal, and the off-diagonal represents the common SNPs deriving from pairs comparisons among breeds. PON had the highest number of monomorphic SNPs (n = 4,151), while they amounted to 3,456 for CHA and 3,915 for LIM. MUP and SAB presented numbers close to 2,000 while SAM and SAR had the lowest values of monomorphic SNPs (1,663 and 1,447, respectively). As expected, the two breeds under selection shared the greatest number of monomorphic SNPs and were followed by LIM vs. PON and PON vs. CAL. However, no monomorphic SNP was found in common in all eight breeds.

**Fig 1.**
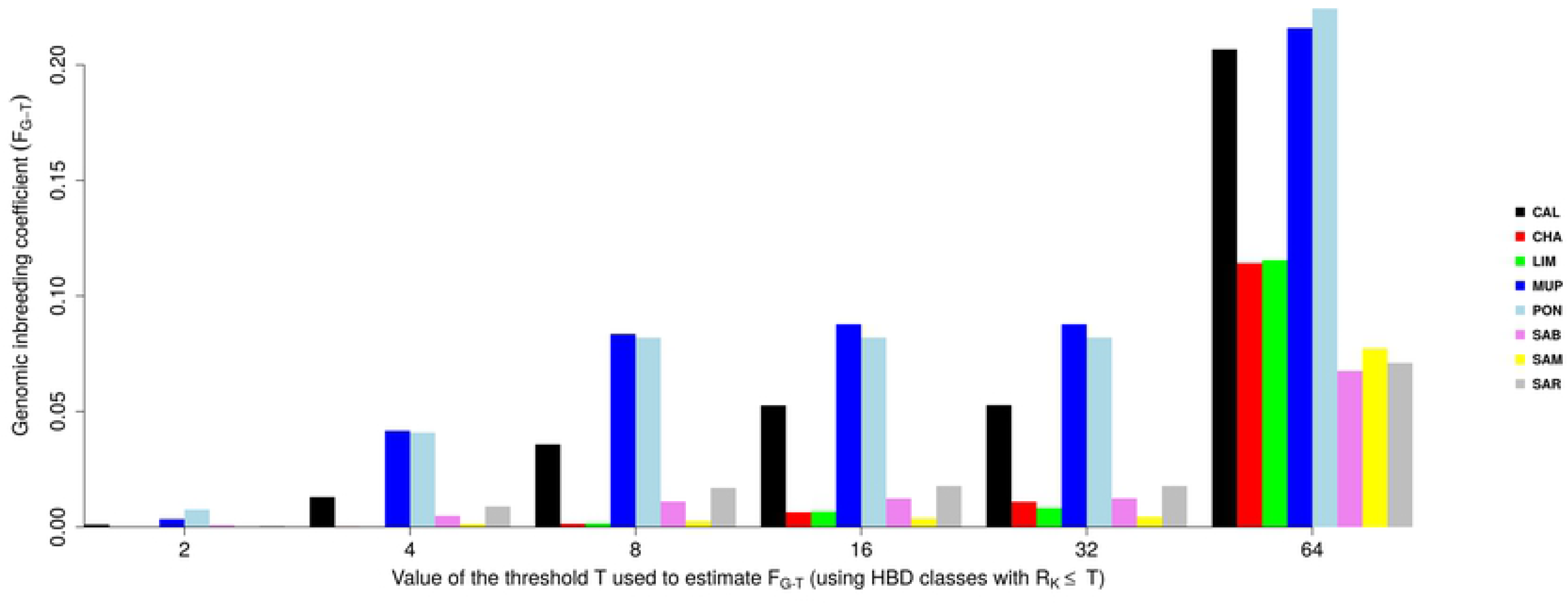
Heatmap of monomorphic SNPs pairs comparison among breeds; CAL = Calvana; CHA = Charolais; LIM = Limousine; MUP = Mucca Pisana; PON = Pontremolese; SAB = Sardo Bruna; SAM = Sardo Modicana; SAR = Sarda.

The monomorphic SNPs distribution across autosomes has been investigated and it is showed in Fig 2. As expected, the amounts of monomorphic SNPs decreased relative to chromosomic length. However, the highest total number was reported in BTA5, except for SAR and SAM (BTA6); the lowest number of monomorphic SNPs was found in BTA25 for Tuscan breeds and LIM, in BTA26 for Sardinian breeds and lastly, in BTA28 for CHA.

**Fig 2.**
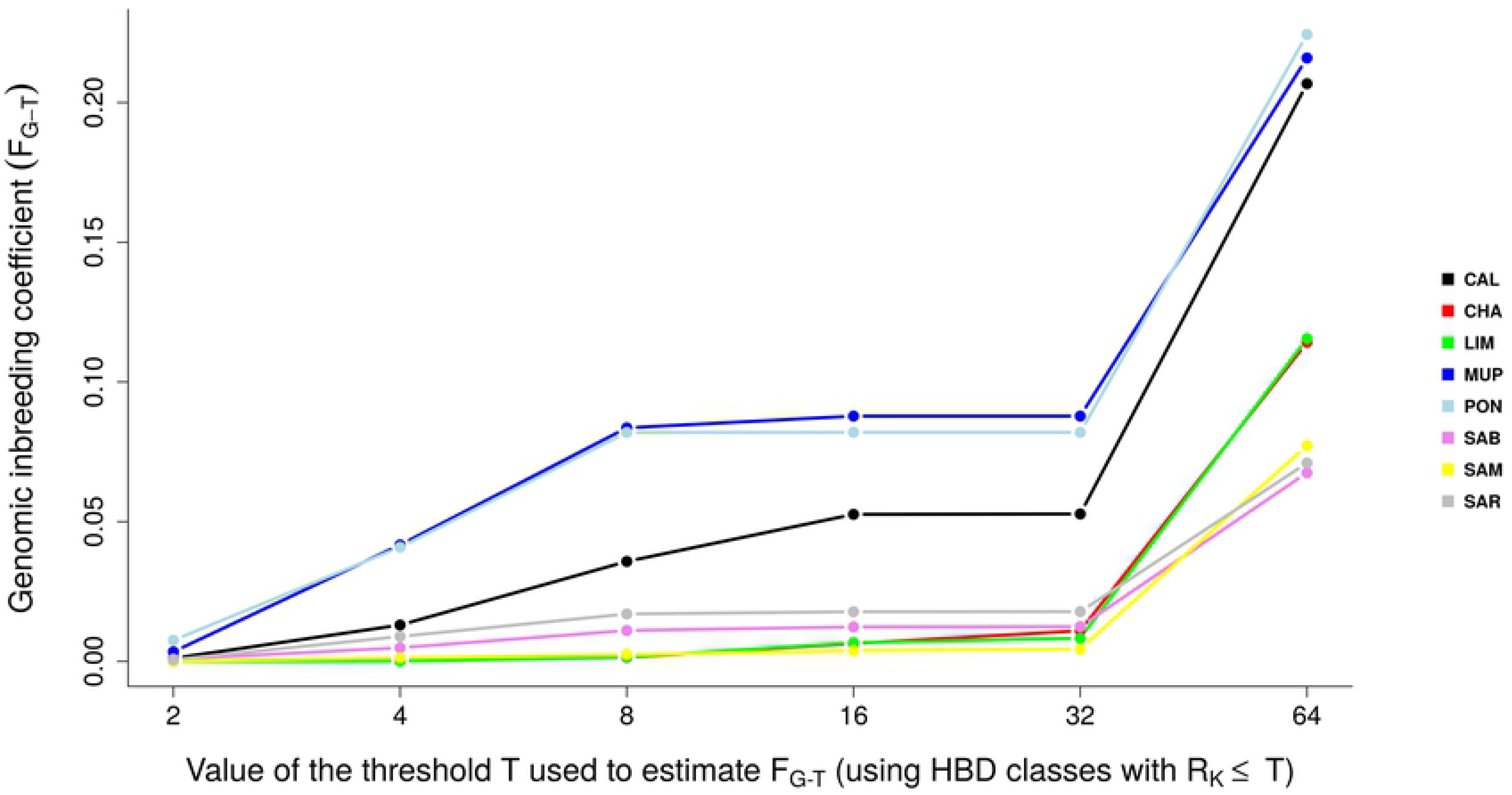
Monomorphic Single Nucleotide Polymorphisms distribution across chromosomes in each breed^1^, where ^1^ CAL = Calvana; CHA = Charolais; LIM = Limousine; MUP = Mucca Pisana; PON = Pontremolese; SAB = Sardo Bruna; SAM = Sardo Modicana; SAR = Sarda.

### Multidimensional scaling plot analysis

The MDS plot evidenced clustering of breeds (Figure 3).

**Fig 3.**
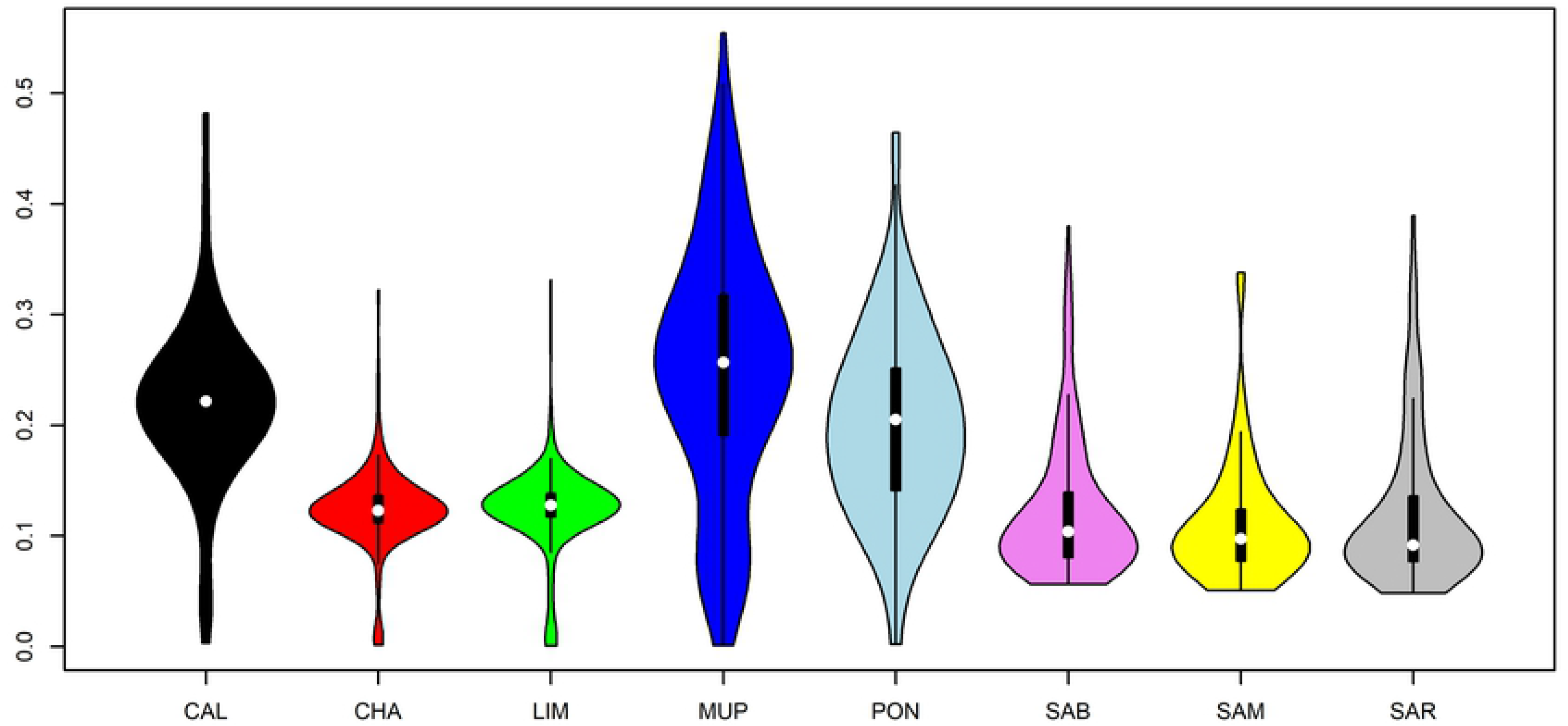
Multidimensional scaling plot for 8 cattle beef breeds, where CAL = Calvana; CHA = Charolais; LIM = Limousine; MUP = Mucca Pisana; PON = Pontremolese; SAB = Sardo Bruna; SAM = Sardo Modicana; SAR = Sarda.

MUP and CAL were extremely distant from each other and from the other six breeds, which grouped in a unique large cluster. Only a small group of MUP samples were nearer to the third cluster (Sardinian and Cosmopolitan breeds). Both LIM and CHA showed extremely compact clusters, suggesting as expected, a low genetic variability within each breed, and also close to each other, underlining their different genetic background compared to local breeds. SAM and SAB individuals were more scattered than SAR and PON which, however, overlapped with these former.

### Runs of Homozygosity detection

Table 2 shows the total number of ROH detected per breed, the average number per individual and the ROH total length.

**Table 2.**
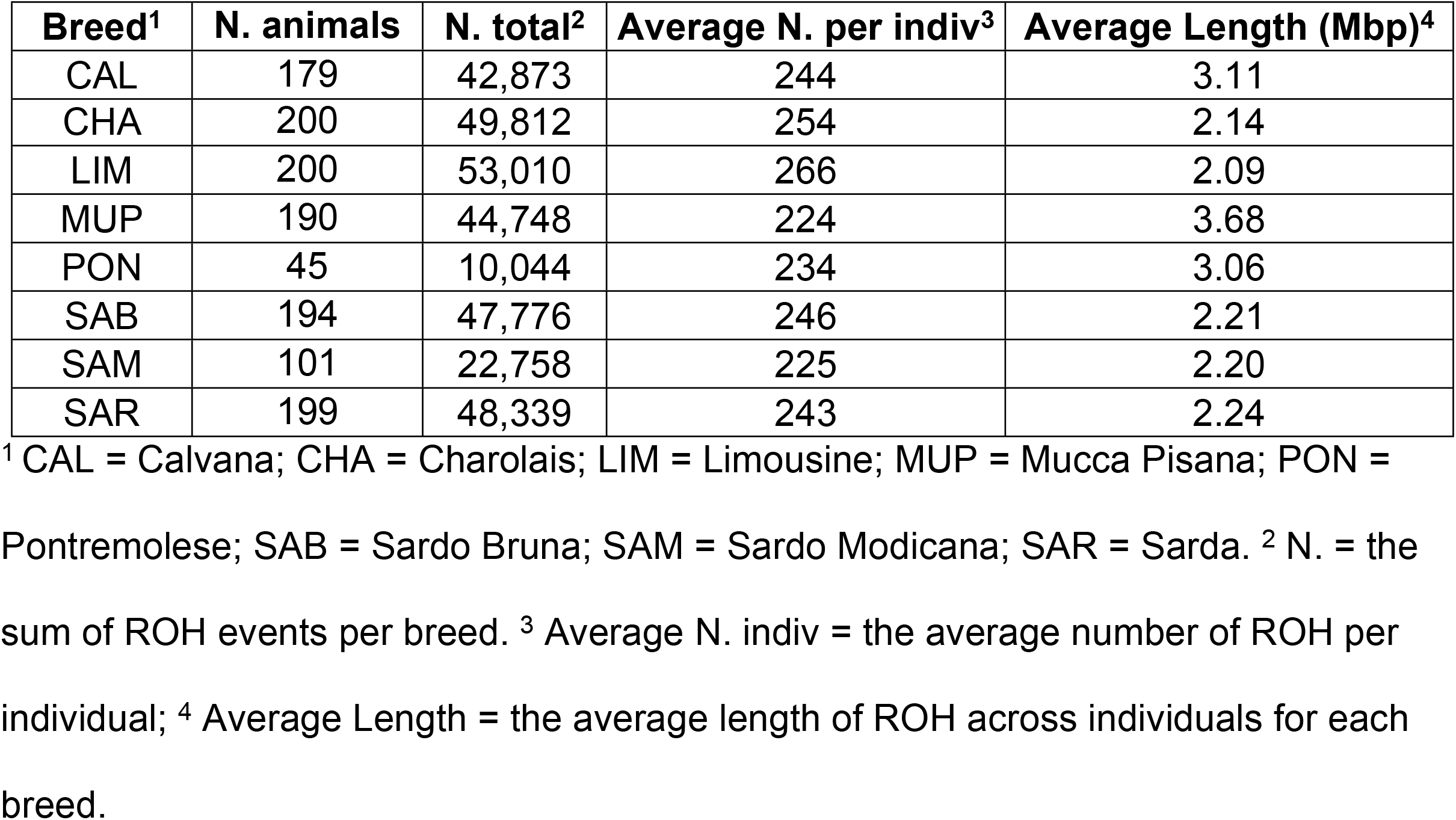
Descriptive statistics of ROH for the eight breeds.

| Breed<sup>1</sup> | N. animals | N. total<sup>2</sup> | Average N. per indiv<sup>3</sup> | Average Length (Mbp)<sup>4</sup> |
| --- | --- | --- | --- | --- |
| CAL | 179 | 42,873 | 244 | 3.11 |
| CHA | 200 | 49,812 | 254 | 2.14 |
| LIM | 200 | 53,010 | 266 | 2.09 |
| MUP | 190 | 44,748 | 224 | 3.68 |
| PON | 45 | 10,044 | 234 | 3.06 |
| SAB | 194 | 47,776 | 246 | 2.21 |
| SAM | 101 | 22,758 | 225 | 2.20 |
| SAR | 199 | 48,339 | 243 | 2.24 |

<sup>1</sup> CAL = Calvana; CHA = Charolais; LIM = Limousine; MUP = Mucca Pisana; PON =
Pontremolese; SAB = Sardo Bruna; SAM = Sardo Modicana; SAR = Sarda. <sup>2</sup> N. = the
sum of ROH events per breed. <sup>3</sup> Average N. indiv = the average number of ROH per
individual; <sup>4</sup> Average Length = the average length of ROH across individuals for each
breed.

The highest number of ROH identified were found in the two cosmopolitan breeds (LIM=53,010; CHA= 49,812), whereas SAM and PON showed the lowest values, 22,758 and 10,044, respectively. However, the average number per individuals was almost the same in each breed, ranging from 224 in SAB to 266 in SAR. For each chromosome the number of ROH per breed (S1 Table) has been calculated. *Bos taurus* autosome (BTA) 1 had consistently the highest ROH number in all breeds, except for MUP, where BTA2 had the highest number of ROH. The average length was found to be higher in Tuscan breeds (ranging from 3.06 to 3.68 Mbp). The Sardinian breeds had intermediate values (2.20 – 2.24 Mbp) while the average ROH length for the cosmopolitan breeds was 2.14 Mbp for CHA and 2.09 Mbp for LIM. PCA analysis on the number of ROH per chromosome revealed groups among breeds (Fig 4). PC1 clearly separated Tuscan breeds from Sardinian and cosmopolitan, while PC2 placed SAR, SAB, CHA and LIM close to 0, while MUP and CAL located to the opposite site to PON. The first two PCs explained together the 95.26% of the total variation among samples.

**Fig 4.**
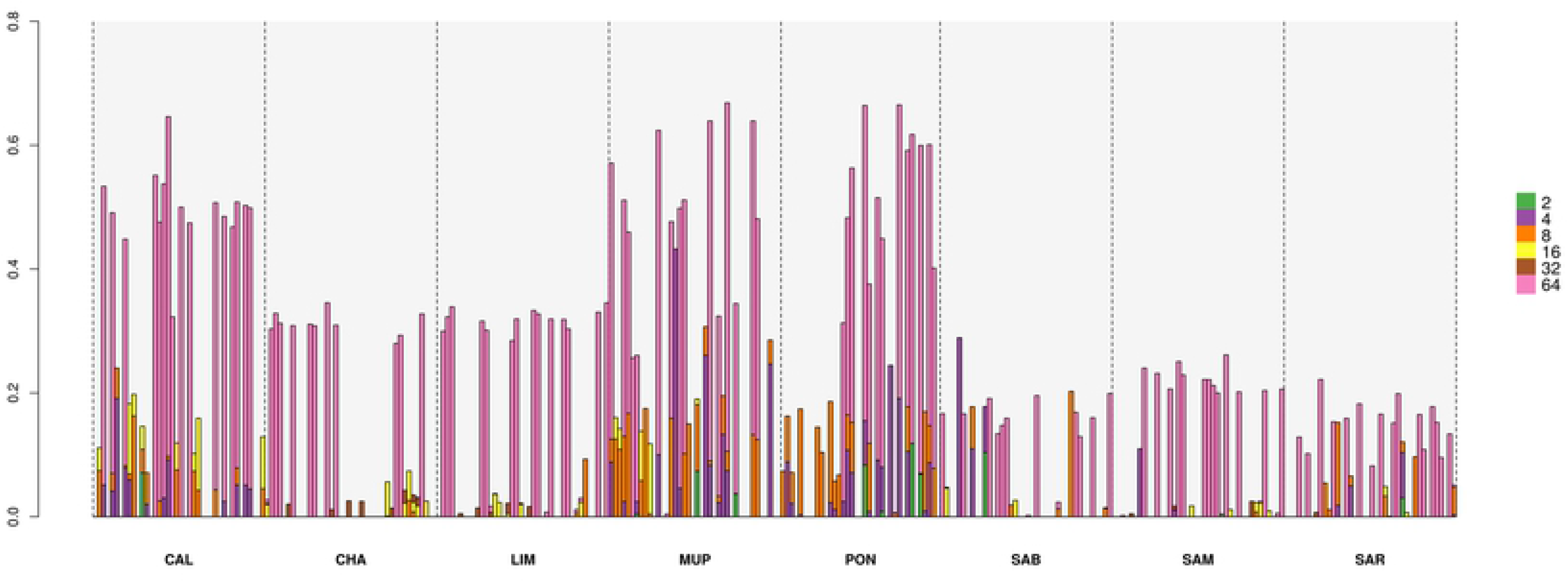
Scatterplot of the first two principal components (PCs). Principal component analysis (PCA) was performed on the number of identified ROH per chromosome in each breed. CAL = Calvana; CHA = Charolais; LIM = Limousine; MUP = Mucca Pisana; PON = Pontremolese; SAB = Sardo Bruna; SAM = Sardo Modicana; SAR = Sarda.

Five classes of length were considered in order to investigate the ROH pattern. The ROH distribution by length and number for each breed were reported in Fig 5.

**Fig 5.**
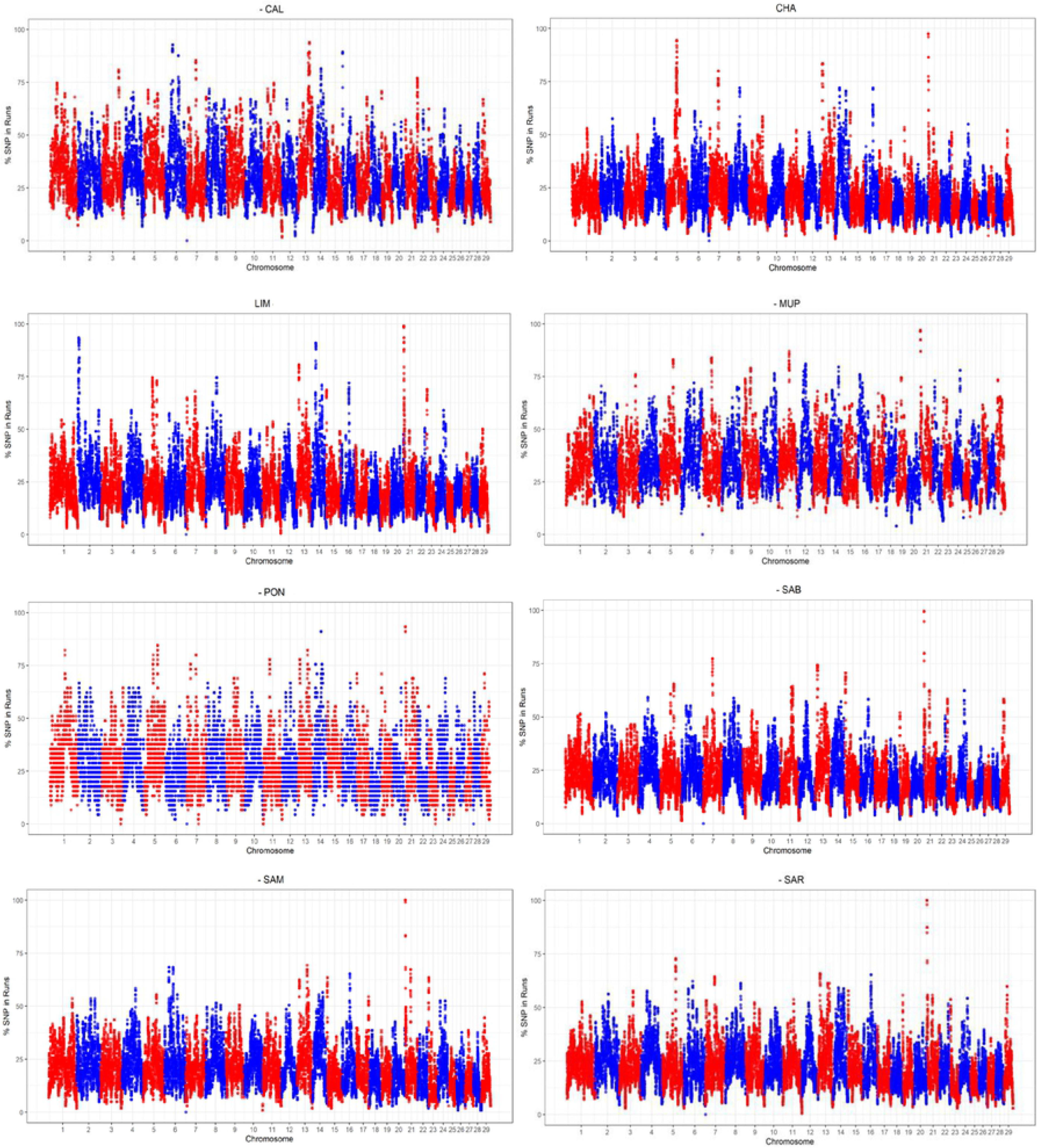
ROH classified into 5 classes of length. CAL = Calvana; CHA = Charolais; LIM = Limousine; MUP = Mucca Pisana; PON = Pontremolese; SAB = Sardo Bruna; SAM = Sardo Modicana; SAR = Sarda.

The majority of ROH detected belonged to the first two classes (<2 and 2-4 Mbp) for all breeds. LIM and SAR had ∼34,000 ROH with length less than 2 Mbp, followed by CHA and SAB with ∼32,000. PON and SAM were the two breeds with a lower number of short ROH detected (6,200 and 15,602, respectively). The second class of ROH length (2-4 Mbp) maintained similar pattern, with LIM, CHA, SAB and SAR having the higher number of ROH. Regarding the classes of longer length (>4Mbp), CAL and MUP had the highest number of ROH, even in the class of >16 Mbp (1,943 for MUP and 1,018 for CAL). The other six breeds in this last class showed 510, 392, 306, 169, 146 and 108 long ROH, for SAR, SAB, PON, SAM, CHA and LIM, respectively.

### Genomic inbreeding (F_ROH_)

Descriptive statistics for F_ROH_ were reported in Fig 6.

**Fig 6.**
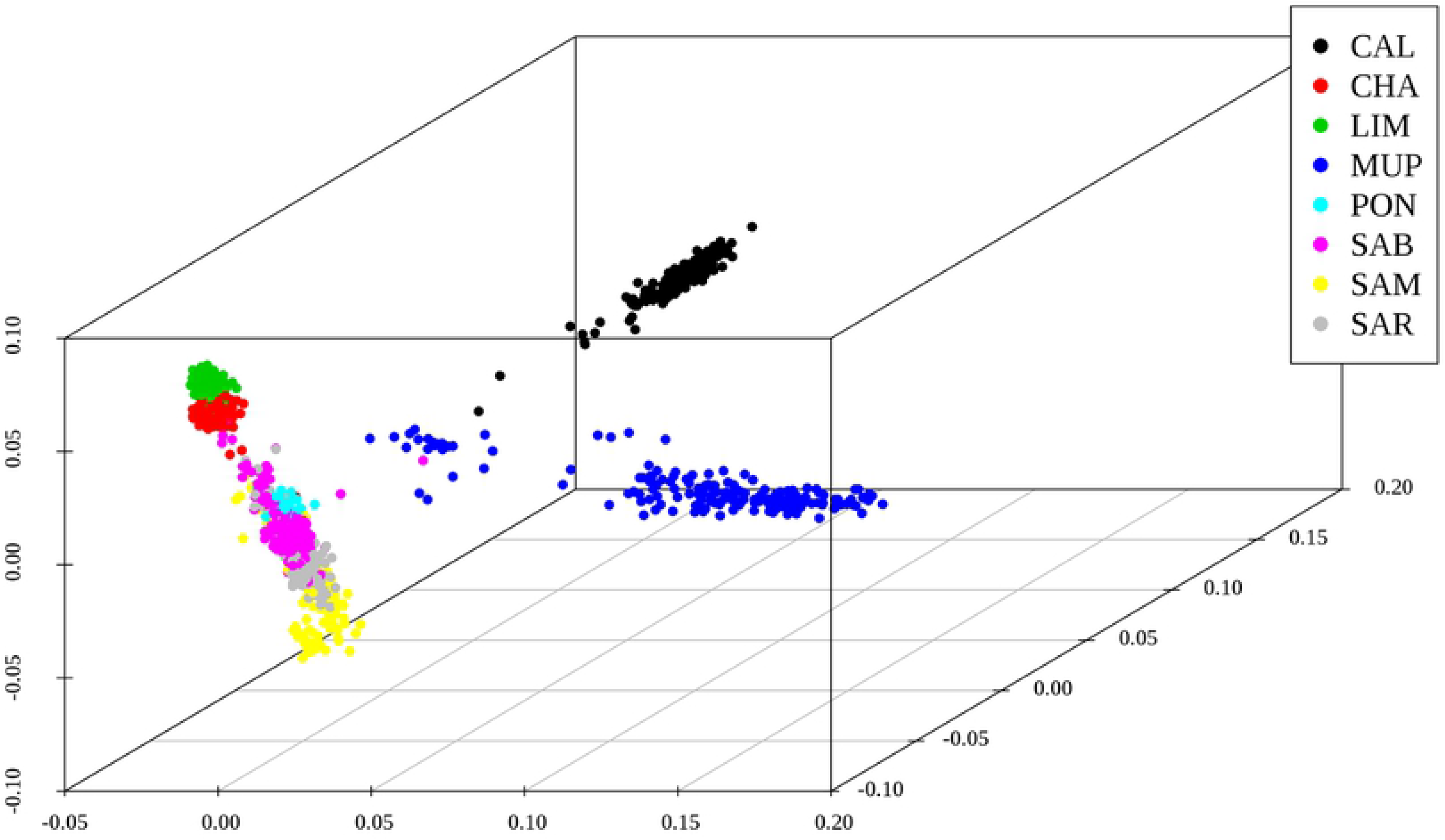
Descriptive statistics of FROH distribution for the eight breeds. CAL = Calvana; CHA = Charolais; LIM = Limousine; MUP = Mucca Pisana; PON = Pontremolese; SAB = Sardo Bruna; SAM = Sardo Modicana; SAR = Sarda.

Tuscan breeds presented the highest level of average genomic inbreeding coefficients (0.33, 0.30, 0.28 for MUP, CAL and PON, respectively) with maximum values that exceeded 0.5. F_ROH_ for LIM, CHA, SAB and SAR was identical (∼0.22), while the lowest average inbreeding coefficient was found in SAM (0.20). Tuscan and cosmopolitan breeds presented minimum values close to 0, instead Sardinian breeds had a minimum F_ROH_ near to 0.14.

To investigate the recent and past inbreeding in each breed, F_ROH_ was calculated for the previous classes of length (Table 3).

**Table 3.** Mean and standard deviation (SD) of F_ROH_ per class of length for each breed.

|  | <2 Mbp |  | 2 – 4 Mbp |  | 4 – 8 Mbp |  | 8 – 16 Mbp |  | >16 Mbp |  |
| --- | --- | --- | --- | --- | --- | --- | --- | --- | --- | --- |
| **Breed<sup>1</sup>** | Mean | SD | Mean | SD | Mean | SD | Mean | SD | Mean | SD |
| CAL | 0.30 | 0.07 | 0.22 | 0.08 | 0.15 | 0.07 | 0.11 | 0.07 | 0.06 | 0.06 |
| CHA | 0.22 | 0.05 | 0.12 | 0.03 | 0.04 | 0.03 | 0.02 | 0.02 | 0.02 | 0.02 |
| LIM | 0.22 | 0.06 | 0.12 | 0.04 | 0.04 | 0.03 | 0.02 | 0.02 | 0.02 | 0.03 |
| MUP | 0.33 | 0.11 | 0.25 | 0.12 | 0.21 | 0.11 | 0.18 | 0.10 | 0.13 | 0.08 |
| PON | 0.29 | 0.08 | 0.20 | 0.08 | 0.14 | 0.09 | 0.12 | 0.08 | 0.09 | 0.07 |
| SAB | 0.22 | 0.06 | 0.12 | 0.06 | 0.05 | 0.06 | 0.05 | 0.06 | 0.05 | 0.05 |
| SAM | 0.20 | 0.05 | 0.10 | 0.06 | 0.05 | 0.06 | 0.04 | 0.05 | 0.04 | 0.05 |
| SAR | 0.22 | 0.06 | 0.12 | 0.07 | 0.05 | 0.07 | 0.06 | 0.07 | 0.06 | 0.06 |

<sup>1</sup> CAL = Calvana; CHA = Charolais; LIM = Limousine; MUP = Mucca Pisana; PON = Pontremolese; SAB = Sardo Bruna; SAM = Sardo Modicana; SAR = Sarda

F_ROH_ in Tuscan breeds was higher for all the classes analyzed when compared to Sardinian and cosmopolitan breeds. The mean for the first class corresponded to the general average inbreeding for each breed, but values decreased with an increasing length. Sardinian breeds maintained values near to 0.05 from 4-8 Mbp up to >16 Mbp classes while in the latter length class, CHA and LIM inbreeding coefficients were close to 0. MUP showed the highest F_ROH_ in all the 5 categories, having an average genomic inbreeding coefficient of 0.13 in long ROH.

### Selection signatures and Gene enrichment

To identify genomic regions potentially important for selection and/or conservation, the SNPs’ frequency contained in the runs were plotted across autosomes for each breed (Fig 7). As presented in Fig 7, the autosomes generally more interested by ROH with high occurrence was BTA21, except for CAL, which were BTA13 and BTA6. LIM and CHA presented ROH peaks also on BTA2 and BTA5, respectively.

**Fig 7.**
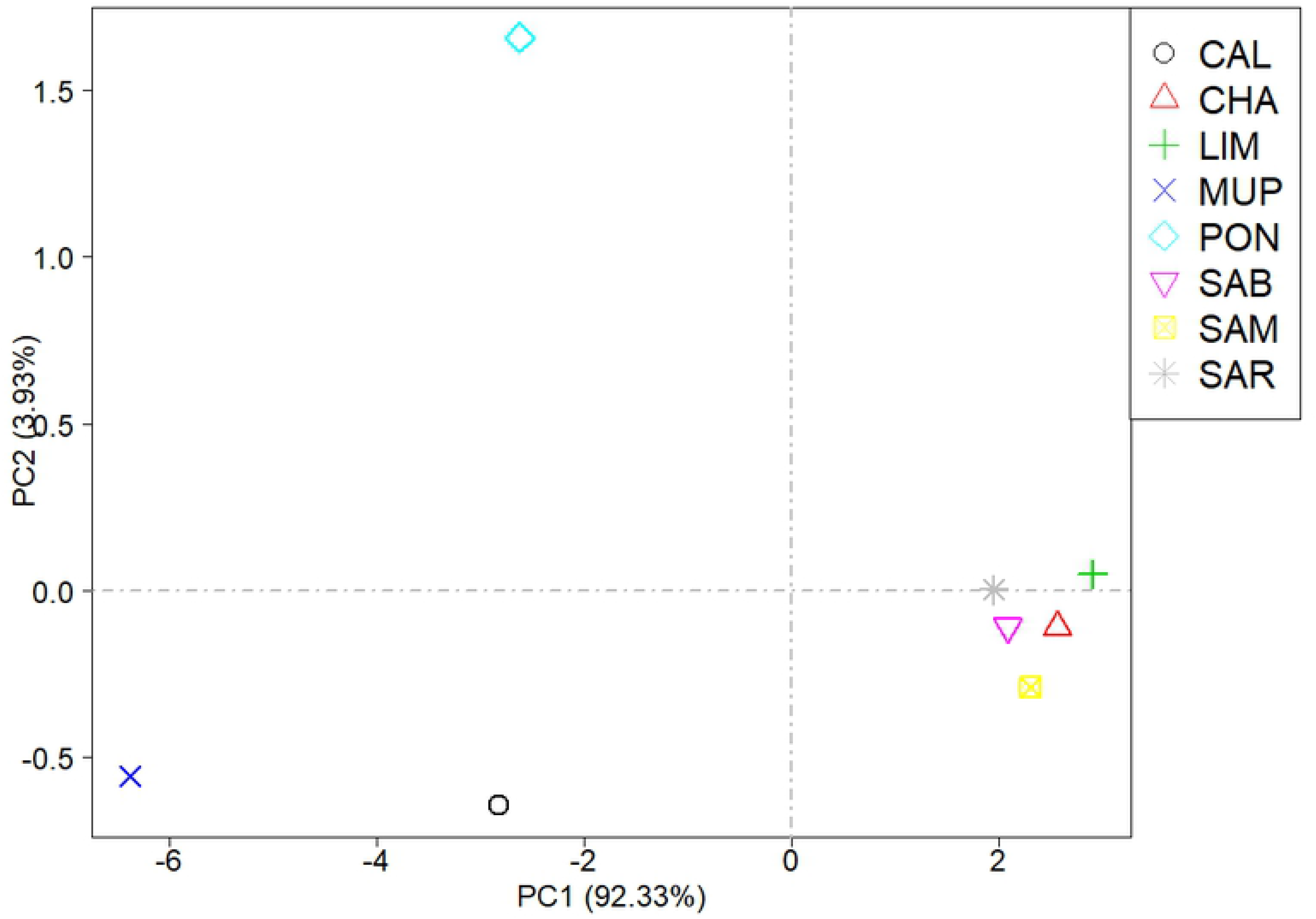
Manhattan plots of the distribution of ROH in the eight cattle breeds. The x-axis is the SNP position and the y-axis shows the frequency (%) at which each SNP was observed in ROH across individuals (CAL = Calvana; CHA = Charolais; LIM = Limousine; MUP = Mucca Pisana; PON = Pontremolese; SAB = Sardo Bruna; SAM = Sardo Modicana; SAR = Sarda).

Applying the abovementioned threshold of 80%, 35 ROH have been detected in total across all breeds (S2 Table); the highest number of genomic regions identified was found in Tuscan breeds (n=11, CAL; n= 9, MUP; n= 6, PON). Five ROH with high occurrence were found both for CHA and LIM, while each Sardinian breed presented only one run. The longest runs were found in LIM (2.65 and 2.38 Mbp), followed by MUP (2.17 Mbp), SAM and SAR (2 Mbp).

One genomic region on BTA21 was found in common in several breeds starting at 83,766 Mbp with BovineHD2100000012. The region was almost identical for CHA, PON, SAB with a length of ∼1.70 Mbp; this run contained 18 SNPs for CHA and PON, 17 for SAB. This ROH was present also in LIM, MUP, SAM, and SAR starting at the same SNP but finishing with different markers (BovineHD2100000320 for LIM located at 2,467,774 bp, BovineHD2100000283 for MUP at 2,256,102 bp and BovineHD2100000258 for SAM and SAR positioned at 2,085,345 bp). For the aforementioned breeds, the number of SNPs within this region ranged from 25 (LIM) to 20 (Sardinian). Within this shared run, four genes were detected: *IGHM* (Immunoglobulin heavy constant Mu), *MKRN3* (Makorin finger protein 3), *MAGEL2* (MAGE family member L2) and *NDN* (Necdin), located upstream of the Prader-Willi syndrome (PWS) region.

From the 35 ROH detected, within-breed specific regions containing a minimum number of 15 SNPs were selected to investigate the genes (Table 4). The list of genes in each run is reported in S3 Table.

**Table 4.** Characterization of within-breed common runs of homozygosity at a threshold of 80% and with at least 15 SNPs and the number of genes included.

| Breed<sup>1</sup> | CHR | Start_SNP | End_SNP | Length_Mbp | N. genes |
| --- | --- | --- | --- | --- | --- |
| CAL | 6 | BovineHD0600010715 | Hapmap26233-BTA-75846 | 1.49 | 2 |
|  | 16 | BovineHD1600000011 | BovineHD1600000286 | 1.06 | 11 |
| CHA | 5 | BovineHD0500016070 | BovineHD0500016088 | 0.09 | 3 |
|  | 5 | BovineHD0500016090 | BovineHD0500016469 | 1.74 | 57 |
| LIM | 2 | ARS-BFGL-NGS-21306 | BTB-01111412 | 2.65 | 22 |
|  | 14 | BovineHD1400007190 | Hapmap40958-BTA-34312 | 1.70 | 7 |
| MUP | 5 | 5_74951342 | 5_75130860 | 0.18 | 1 |
| PON | 5 | BovineHD0500021258 | 5_75130860 | 0.37 | 6 |

<sup>1</sup> CAL = Calvana; CHA = Charolais; LIM = Limousine; MUP = Mucca Pisana; PON = Pontremolese.

Among genes presented in S3 Table, we found some of special interest. For CAL, we found the *MYOG* (Myogenin) (a muscle-specific transcription factor), *CHI3L1* (Chitinase 3 Like 1) and *CHIT1* (Chitinase 1) genes on BTA16, both of these are involved in inflammatory processes. On BTA5, for CHA we found the largest number of genes (n = 57). These were linked to cell survival after damage or stress (*TIMELESS*; Timeless Circadian Regulator), transport and/or esterification of cholesterol (*APOF*; Apolipoprotein F), growth regulation, development and differentiation (*SLC39A5* - Solute Carrier Family 39 Member 5, *PA2G4* - Proliferation-Associated 2G4, *CD63* - CD63 Molecule) and olfactory receptors (*OR10P1*, *OR6C4*, *OR2AP1*, *OR6C2*, *OR6C68*). For LIM, the most interesting detected genes were located on BTA2: *CYP27C1* (encoding a member of the cytochrome P450 superfamily), *MSTN* (Myostatin) and genes encoding collagen chains (*COL5A2* - Collagen Type V Alpha 2 Chain, *COL3A1* - Collagen Type III Alpha 1 Chain). MUP and PON shared on BTA5, *CACNG2* (Calcium Voltage-Gated Channel Auxiliary Subunit Gamma 2), which is a gene involved in synaptic plasticity, learning and memory.

### Homozygosity by descent (HBD) segments and global inbreeding

The percentage of non-HBD segments was higher in SAB (42.7%), followed by SAM, SAR and MUP (39%), while CHA, LIM and PON showed values around 32.5%. The HBD segments identified (Fig 8) belonged mainly to HBD class with R_K_ equal to 64. Tuscan breeds showed a greater proportion of HBD genome also for R_K_ ranged from 4 to 8, while CAL is the breed with higher proportion of HBD segments when R_K_ was 16.

**Fig 8.**
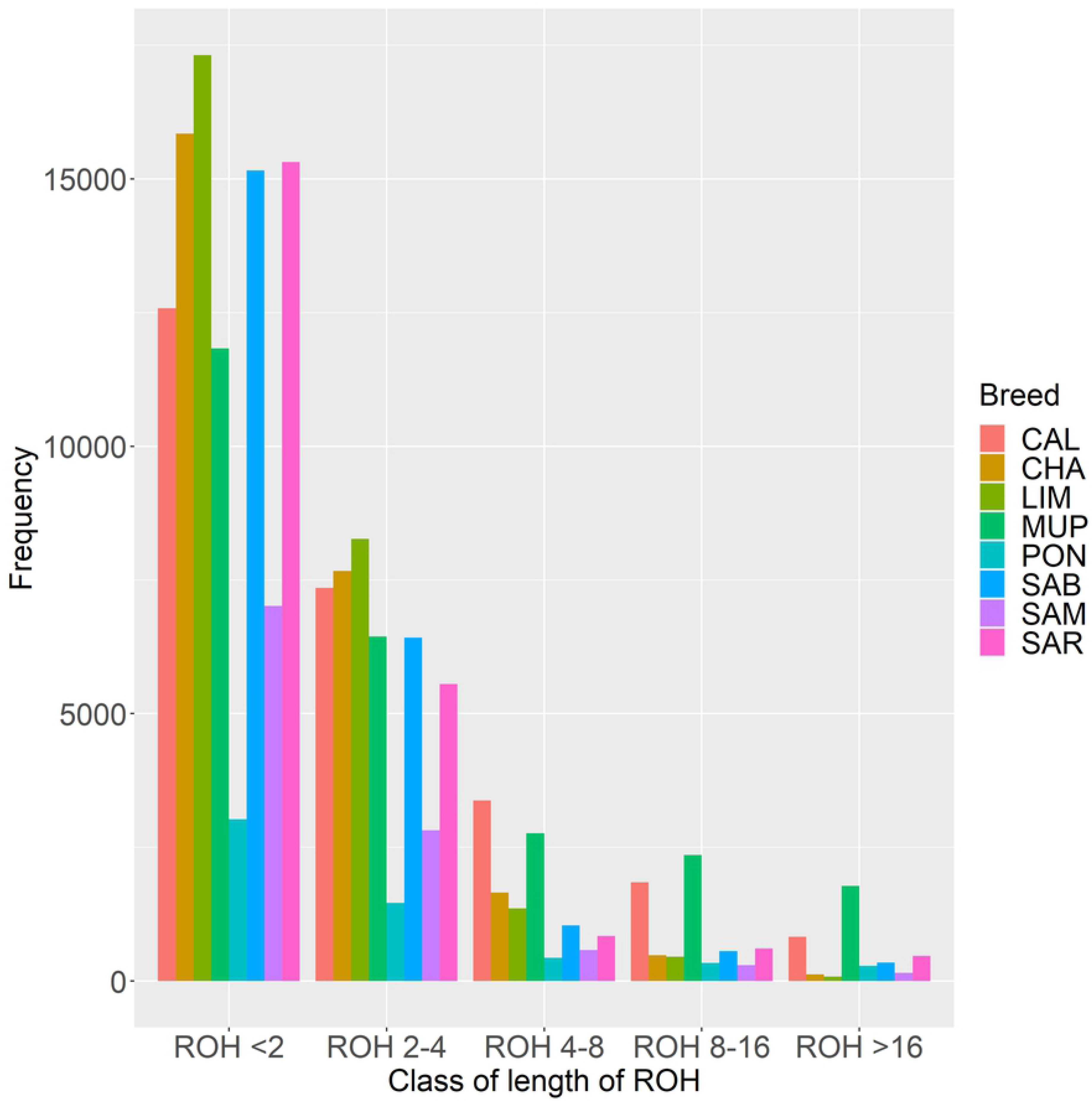
Proportion of the genome consisted of HBD classes in different RK. CAL = Calvana; CHA = Charolais; LIM = Limousine; MUP = Mucca Pisana; PON = Pontremolese; SAB = Sardo Bruna; SAM = Sardo Modicana; SAR = Sarda.

The results suggested that all the breeds suffered an increase in inbreeding during ancient generations (around 32 generations ago), and only Tuscan breeds have been involved in new consistent inbreeding events, approximately 2-4 generations ago. Results were similar when inbreeding (F_G-T_) was calculated respect to different base populations (Fig 9). The F_G-T_ was estimated as the probability of belonging to any of the HBD classes with a R_K_ ≤ a threshold T, averaged over the whole genome.

**Fig 9.**
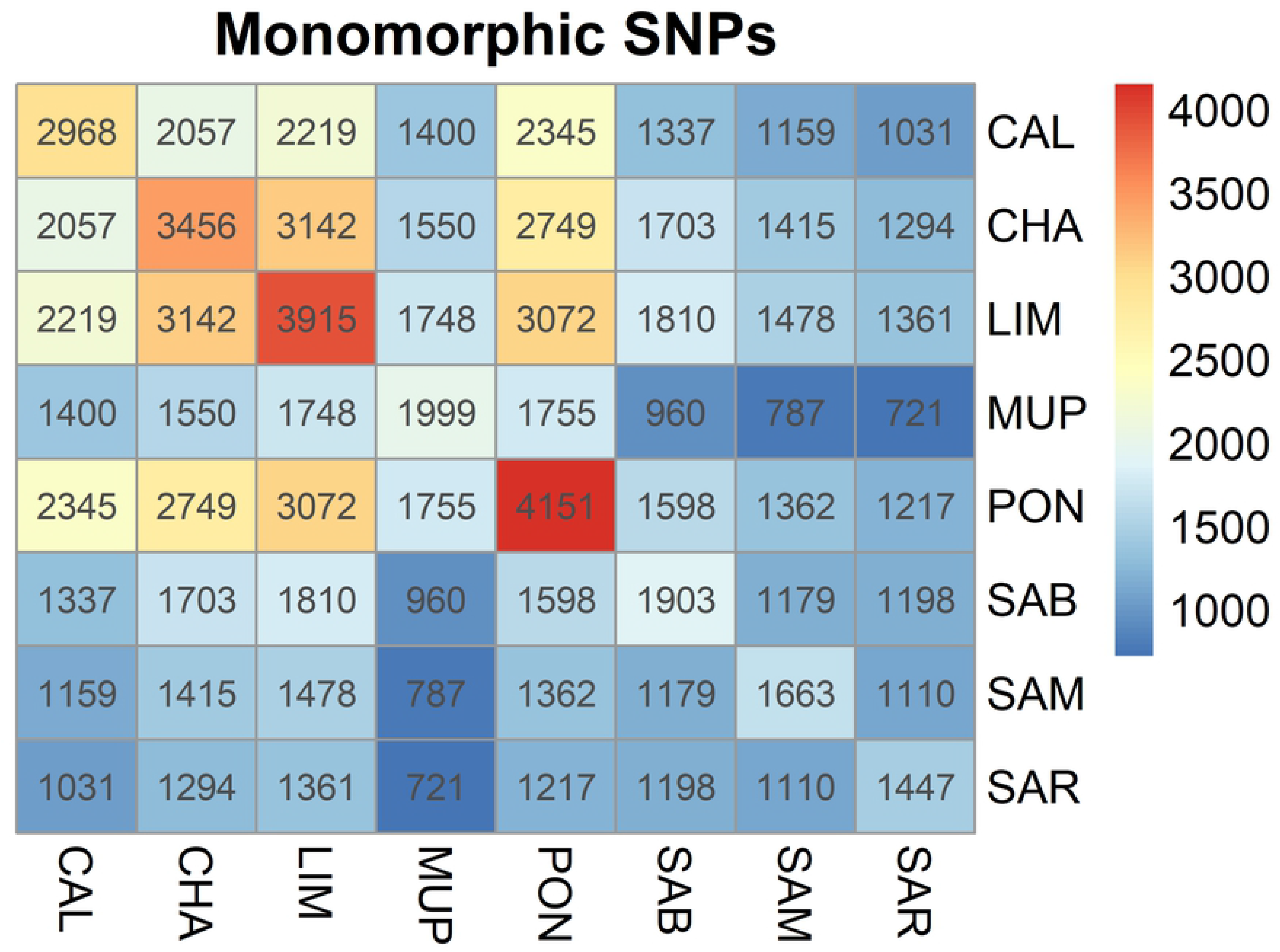
Genomic inbreeding coefficients estimated respect to different base populations (F_G-T_) selecting different thresholds T, setting the base population approximately 0.5 * T generations ago.

The F_G_ estimated with the most remote base population showed values exceeding 0.2 for Tuscan breeds, while LIM and CHA had F_G_ around 0.1. Sardinian maintained F_G_ close to 0.06. Classes associated with smaller R_K_ rates (i.e., with longer HBD segments) explained a smaller HBD proportion: the average inbreeding coefficient was close to 0 in cosmopolitan and Sardinian breeds, while for PON and MUP F_G_ was slightly lower than 0.1 from classes with 8 ≤ R_K_ ≤ 32, while CAL didn’t exceed 0.05. The inbreeding coefficient associated with common ancestors tracing back up to approximately two generations ago (corresponding to HBD-classes with R_K_ ≤ 4) tended to 0 in all Tuscan breeds.

Partitioning of individual genomes in different HBD classes was also performed and Fig 10 reports the plot of 40 randomly chosen individuals per breed. Each bar represents an individual, the white spaces are individuals with no HBD segments belonging to HBD classes analyzed, the height of the different stacks is the proportion of the genome associated with the HBD class of the corresponding color and the total height showed the overall level of inbreeding.

**Fig 10.**
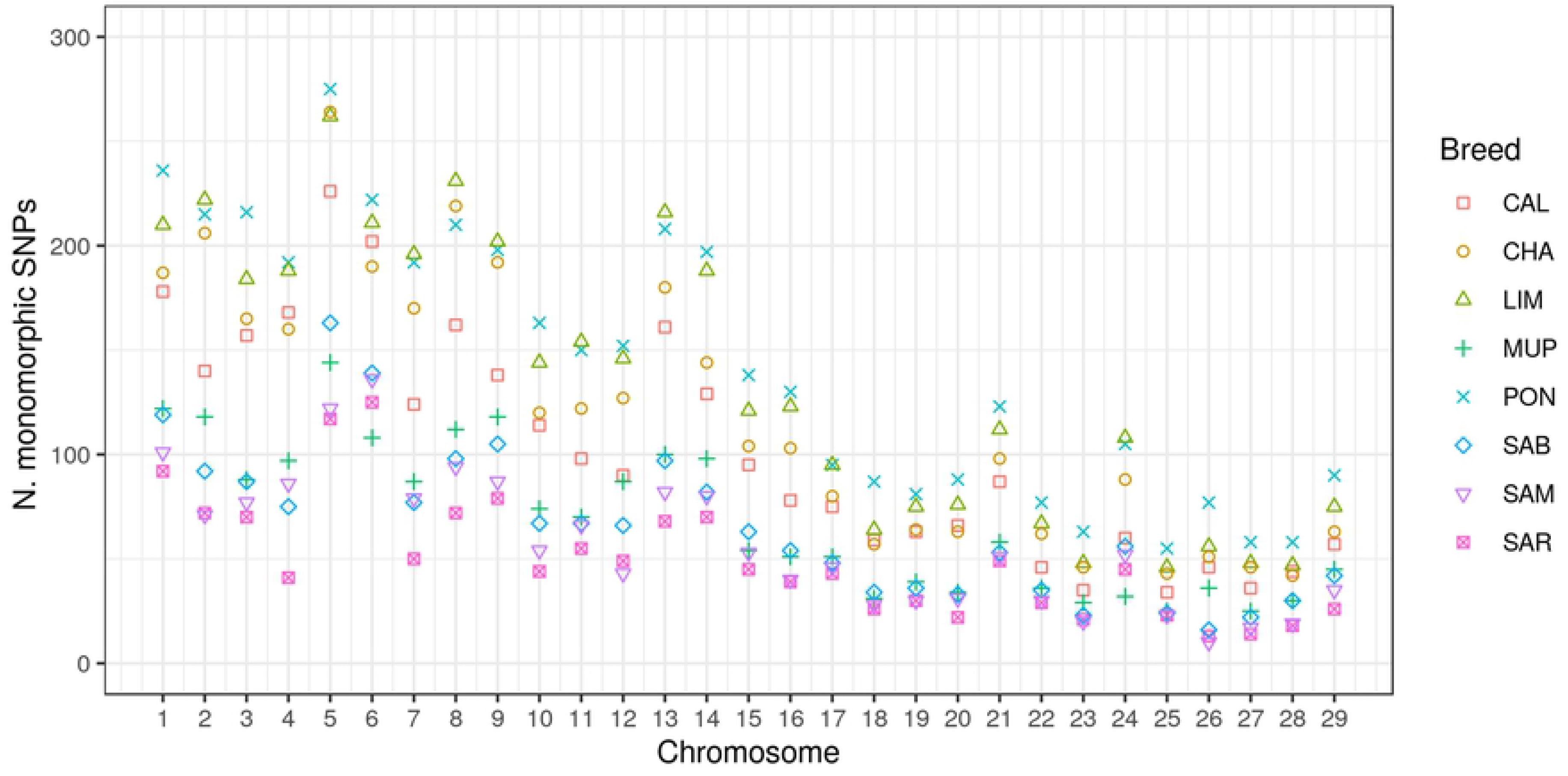
Proportion of the genome interested by HBD classes in different R_K_. CAL = Calvana; CHA = Charolais; LIM = Limousine; MUP = Mucca Pisana; PON = Pontremolese; SAB = Sardo Bruna; SAM = Sardo Modicana; SAR = Sarda.

The results confirmed that all breeds acquired the majority of their inbreeding load derived from ancient ancestors (32 generations ago), but once again, the Tuscan breeds appeared with a different demographic historical structure compared to Sardinian and cosmopolitan breeds; the level of inbreeding was higher in Tuscan and deriving both from past and recent phenomena. Indeed, PON and MUP were affected also by inbreeding acquired in recent generations (2-4, i.e. R_K_=4 and 8, respectively). CAL also showed traces of ancestors from 8 generation ago in several individuals. Sardinian and cosmopolitan breeds showed lower levels of inbreeding and HBD segments derived from ancient ancestors.

## Discussion

The advent of high-throughput genotyping arrays facilitated the study of genetic diversity and population structure in cattle [21], but local breeds remained understudied, even if in the last years greater importance has been given to the maintenance of biodiversity and the adoption of conservation measures for breeds at risk of extinction. Several advantages are brought with the conservation of local breeds, such as economical and genetic benefits [34]. This study comprehensively describes the genome-wide autozygosity and the consequent population structure of six local breed reared in Italy, namely Calvana, Mucca Pisana, Pontremolese, Sardo Bruna, Sardo Modicana and Sarda, by exploring the distribution of ROH, the level of genomic inbreeding (F_ROH_) and the partitioning of homozygous identical by descent (HBD) segments across generations.

### Multidimensional scaling plot analysis

MDS approach has been preferred to Principal component analysis (PCA) because it detects meaningful dimensions that explain observed genetic distance, i.e. pairwise IBS distance, while PCA method calculates the population structure based on genetic correlations among individuals [28]. The genome feature analysis carried out using MDS decomposition (Fig 3) was in accordance with the ROH-based PCA (Fig 4), highlighting a grouping among breeds. In particular, in both analyses Sardinian and cosmopolitan breeds were more similar to each other than Tuscan populations, except for PON, which clustered together with Sardinian and cosmopolitan breeds in MDS but not in PCA. This might be because sampled animals of PON were few due to the little size of population (only 49 living animals are reported [35]), and this could affect PCA results but it does not influence MDS analysis, solely based on genetic distances clustering. Plotting of the eight breeds was therefore a description of the breeds sample size. The SAB and SAR breeds counted approximately 25,000 alive animals [36], Charolais 18,000 and Limousine 50,000 (http://www.anacli.it/), while Calvana and Mucca Pisana samples amounted to a few hundred [34]. CHA and LIM formed the most compact cluster indicating that the breeding management in these breeds has a narrower genetic basis.

### Runs of Homozygosity and Genomic inbreeding (F_ROH_)

The total number of ROH detected in each breed, was higher than what found in other studies focusing on cattle [15,36,37]. Differences might be due to the low-density panel used for genotyping, the quality control of the genotypes, the parameters used to define a ROH and the sample size. The small number of detected ROH in PON and SAM might be a result of limited sample size. The relatively high average length of ROH, ranged from 2.09 to 3.68 (Table 2), suggested that ancient inbreeding is present in all breeds. Short ROH represented the vast majority of ROH detected in all breeds (Fig 5), being more profound for LIM and CHA. This is in line with the history of cosmopolitan breeds, which have seen a crucial increase in sample size in the last 15 years (http://www.anacli.it). The growing interest in selection programs have probably led to a slight increase in inbreeding compared to other European Limousine and Charolais populations; indeed, Szmatola et al. [15] described F_ROH_ in Polish Limousine ranging from 0.059 (>1Mbp) to 0.011 (>16 Mbp), while, Polish Charolaise showed values from 0.065 to 0.009. In this study F_ROH_ decreased in both selected breeds from 0.22 to 0.02 for the aforementioned classes of length.

Nowadays, SAB, SAR and SAM are distributed across 1,432, 950 and 146 farms, and it is known that Sardinian farmers exchange bulls between herds, causing a high average relatedness of individuals within farm but allowing a low degree of kinship among farms [36]. This could explain why F_ROH_ in SAR, SAB and SAM has been maintained near 0.05 (Fig 6 and Table 3) in medium (4-8 Mbp) and long ROH (>16 Mbp), suggesting that ancient and recent inbreeding has created a plateau of consanguinity across Sardinian populations. Results reported by Cesarani et al. [36], where the average length per individual ranged from 2.9 for SAR to 2.4 Mbp for SAM, showed a trend of decreased inbreeding in these populations. Tuscan breeds are in a more worrisome situation with their population sizes and inbreeding levels being at critical levels. The MUP breed presented average F_ROH_ values equal to 0.33 and 0.13 for short and long runs, respectively. The PON and CAL breeds had genomic inbreeding equal to 0.30 in the first mentioned class and 0.9-0.6 in the last, respectively. To the best of our knowledge, studies on Tuscan breeds here investigated were not present in literature, but several studies on local cattle breeds reported lower F_ROH_ values than our results. Addo et al. [38] analyzed two German local cattle populations (Angler and Red-and-White dual-purpose breeds) compared to Red Holstein and, on the contrary of this study, is the cosmopolitan population to have the greater values of genomic inbreeding. However, F_ROH_ decreased quickly in the two local breeds, arriving to 0.009 and 0.02 in >16 Mbp length class, while in the Tuscan breeds F_ROH_ ranged from 0.06 to 0.13.

Quantification of the genome wide autozygosity for genetic conservation aims is fundamental because several studies correlated F_ROH_ with inbreeding depression in production and fertility traits [39–41]. In addition, recent inbreeding could fix recessive deleterious variants because there was a strong positive correlation between the number of deleterious homozygotes and the genomic ROH proportion [42].

### Selection signatures and Gene enrichment

The higher threshold used in this study (80% of frequency) for the investigation of selection signatures, led to the identification of a common run between breeds. It is located on BTA21, starting from 83.766 kbp to 1,786.020 kbp.

An interesting genomic region with an occurrence of more than 80%, has been found in CAL on BTA16 (from 99,900 to 1,163,809 bp), where *MYOG* and *FMOD* genes were located. *MYOG* is related with *MSTN* gene which regulates muscle mass. The different myogenin genotypes are related to the variation in the number of muscle fibers and the growth rate, which lead to a variation in the muscle mass [43]. Indeed, has been suggested to use *MYOG* in marker-assisted selection for improving the growth trait in chicken [44].

*FMOD* plays an important role in the maintenance of mature tissues and has been discovered that reduces scar formation without diminishing the tensile strength in adult wound models (i.e. mice, rats and pig) [45]. It might be related to the higher rusticity and adaptability to harshly areas of local breeds. CHA reported a genomic region dense in genes (Table 4) on BTA5 (from 56,722,571 to 58,464,570 bp). Here two groups of genes are located: the first one included *TIMELESS*, *APON*, *APOF*, STAT2, IL23A and PAN2; the second one contained several genes of Olfactory Receptor Families.

*APOF* and *APON* are apolipoproteins which are component of lipoproteins and it has been showed that overexpression of Apolipoprotein F in mice reduced HDL cholesterol levels by 20-25% by accelerating clearance from the circulation [46].

Olfactory receptors (*ORs*) are essential for mammals to avoid dangers and search food [47]. Nowadays, a few genome-wide association studies reported associations between *ORs* and intake-related traits of livestock. Magalhães et al. [48] argued that olfactory receptors play a role in transferring energy within the cell, participating in the change of *GDP* (guanosine diphosphate) to *GTP* (guanosine triphosphate). Other explanations for the effect of *ORs* on meat traits are their action by promoting the absorption of fatty acids and by differentiating adipocyte, that leads to an increase in accumulation of fat; in addition, their known role to increase the search of food improve the weight gain.

This cluster of genes found in CHA, is in line with artificial selection purposes, as for LIM, which included in the first significant run (BTA2; 5,305,197 - 7,958,492 bp), the presence of *MSTN* gene. It is known that *MSTN* inhibits the proliferation of muscle fibers, regulating muscle mass by negatively influencing cell differentiation via the myogenic regulatory factors (such as *MYOG*) [49]; three traits are associated with this gene: meat color L* (QTL:11644), meat percentage (QTL:11883; QTL:18424) and meat weight (QTL:11694) (https://www.animalgenome.org). Previous studies identified *MSTN* within ROH island in Limousine cattle, highlighting that *MSTN* is a gene under selective pressure for the phenotypic features in Limousine breed, indeed, *MSTN* has a strong positive effect on muscling and it is negative correlated with fat deposition [15, 50].

MUP and PON presented consecutive genomic regions on BTA5, sharing the *CACNG2* gene. Interestingly, this gene was found to be associated with milk protein percentage QTL (https://www.animalgenome.org), which is probably because these two breeds originate from several past crossbreeding events including with Holstein and Schwyz (MUP) [35] and Reggiana (PON) (http://www.anacli.it).

### Homozygosity by descent (HBD) segments and global inbreeding

The parameters used in this analysis were chosen according to ROH results. The length of ROH ranged from >0 and <16 Mbp, considering R_K_ = 2 * generation ago and considering also that the length of the HBD segment is 1/R_K_ Morgans [19], we are interested to investigate HBD classes with R_K_ equal to 2, 4, 8, 16, 32, 64, which correspond to 1, 2, 4, 8, 16 and 32 generations ago. The greater proportion of HBD genome originated from ancient ancestors dates back to 32 generations ago and this is in line with the numerous short ROH detected. An unexpected result was that Sardinian breeds showed almost halved values in HBD classes (when R_K_ is 64) compared to cosmopolitan, suggesting that for these breeds, R_K_ should be increased in order to detect shorter HBD segments which have been found during ROH analysis (Fig 5). The proportion of HBD genome was close to 0 with R_K_ equal to 32 for all breeds and would be interesting to investigate historic events during the 16^th^ generation. No pedigree data on this generation was available when pedigree inbreeding has been calculated in a previous study for these Italian breeds, except for CHA whose results are comparable [34].

However, in the last generations the inbreeding coefficients decreased as described in Fig 9. This suggests that the increased attention to the maintenance of biodiversity have led to a greater mating control by farmers. Unexpectedly, the investigation of individual proportion of HBD genome, identified some individuals that are not HBD. Further analyses are needed but these individuals could be identified and selected for their use in mating programs to decrease inbreeding. Furthermore, animals with a small proportion of HBD genome compared to population could be also useful in conservation plans of local endangered cattle. Nevertheless, Fig 10 underlines the worrisome situation for Tuscan breeds in terms of inbreeding. Also, issues in mating management have been arising since the global inbreeding depends on past ancestors but also to recent generations. Given the limited diffusion of CAL, MUP and PON, the number of potential matings is extremely reduced.

## Conclusion

The genomic results using a low-density SNP chip panel showed critical inbreeding levels in smaller local populations. Cosmopolitan breeds showed lower genetic variability but also negligible inbreeding levels, demonstrating the soundness of the ongoing breeding scheme. The population structure and genetic distances highlighted a clear separation among the breeds, with clusters related to productive purposes and sample sizes. With the present study, we provided a comprehensive overview of two groups of local cattle breeds which are economically and culturally fundamental for the regions where are reared (Tuscany and Sardinia). A conservation and management strategy to increase and/or improve productions for these local breeds should be provided in order to guarantee a greater profit for farmers and, consequently, to overcome the critical situation in population sizes that characterizes the Tuscan local breeds. The results obtained in this study represent a useful tool, proving background information for the correct genetic management and conservation the populations. In addition, the information from genomic analysis plays a crucial role in the development of mating plans, which should be a necessary practice when populations are so small as those analyzed in this study. Low inbreeding is essential for small local breeds to safeguard the native genetic diversity, and so, for preserving biodiversity.

## Acknowledgments

The research was funded by ANACLI through the I-BEEF project PSRN 2014–2020. Sottomisura 10.2: Biodiversità animale. We acknowledge Associazione Nazionale Allevatori delle razze bovine Charolaise e Limousine (ANACLI) for providing the data.

## Supporting information

**S1 Table. Number of ROH per chromosome in each breed, where CAL = Calvana; CHA = Charolais; LIM = Limousine; MUP = Mucca Pisana; PON = Pontremolese; SAB = Sardo Bruna; SAM = Sardo Modicana; SAR = Sarda**

**S2 Table. Characterization of genomic regions with frequency equal to 80% of ROH occurrence, where CAL = Calvana; CHA = Charolais; LIM = Limousine; MUP = Mucca Pisana; PON = Pontremolese; SAB = Sardo Bruna; SAM = Sardo Modicana; SAR = Sarda**

**S3 Table. List of genes within significant ROH (80% of occurrence) with a minimum of 15 SNPs, where CAL = Calvana; CHA = Charolais; LIM = Limousine; MUP = Mucca Pisana; PON = Pontremolese**

